## Supplementary Figures and Tables for "Bacterial chromatin remodeling associated with transcription-induced domains at pathogenicity Islands"

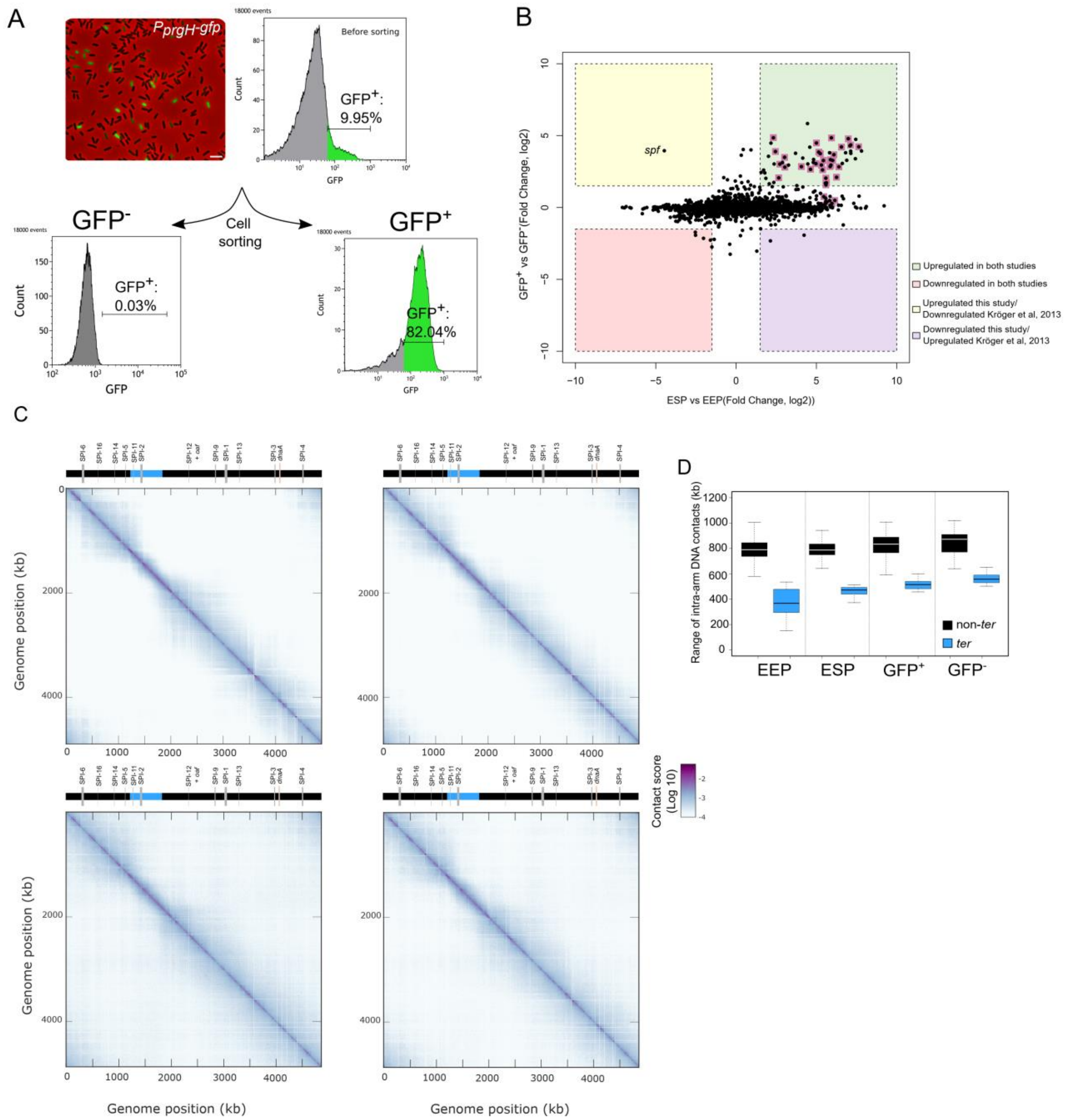


**Supplementary Figure 1. Transcriptomic analyses and chromosome folding in *Salmonella* cells.**

A) Bistable expression of $P_{prgH}$-*gfp* in *S*. Typhimurium as determined by fluorescence microscopy (upper left) or FACS (upper right). Below, expression of $P_{prgH}$-*gfp* after cell sorting in the GFP negative ($GFP^-$) and GFP positive ($GFP^+$) populations.

B) Comparative analysis of genes that are significantly upregulated in this study ($GFP^+$ *vs* $GFP^-$) and in a previous study[1]. SPI-1 genes are highlighted in pink.

C) Normalized contact maps obtained in asynchronous populations of *Salmonella* in early exponentially phase (EEP, upper-left panel), early stationary phase (ESP, upper-right panel), anaerobic growth (lower-left panel) and under oxygen shock (lower right panel).

D) Quantification of the range of intra-arm DNA contacts within *ter* or non-*ter* regions for *S*. Typhimurium Hi-C contact maps. Quantification was performed as previously done in[2]. Boxplot representations are used, indicating the median (horizontal bar), the 25th and the 75th percentile (open box) and the rest of the population except for the outliers (whiskers).

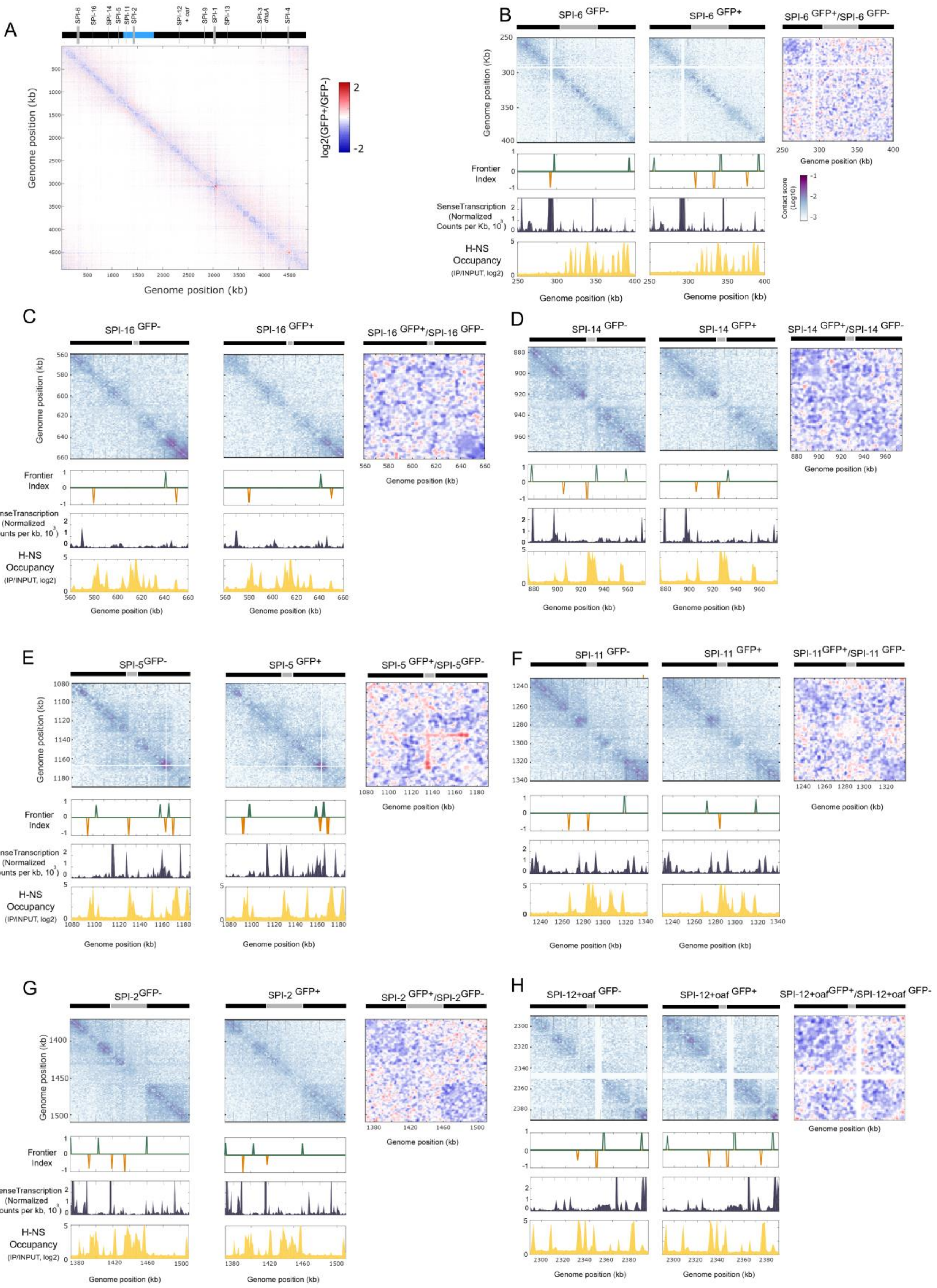
A
SPI-6
SPI-16
SPI-14
SPI-5
SPI-11
SPI-2
SPI-12 + oaf
SPI-9
SPI-1
SPI-13
SPI-3
dnaA
SPI-4
Genome position (kb)
log2(GFP+/GFP-)
B
SPI-6 GFP-
SPI-6 GFP+
SPI-6 GFP+/SPI-6 GFP-
Genome position (Kb)
Frontier Index
SenseTranscription (Normalized Counts per Kb, 10^3)
H-NS Occupancy (IP/INPUT, log2)
Contact score (Log10)
Genome position (kb)
C
SPI-16 GFP-
SPI-16 GFP+
SPI-16 GFP+/SPI-16 GFP-
D
SPI-14 GFP-
SPI-14 GFP+
SPI-14 GFP+/SPI-14 GFP-
E
SPI-5GFP-
SPI-5 GFP+
SPI-5 GFP+/SPI-5GFP-
F
SPI-11 GFP-
SPI-11 GFP+
SPI-11GFP+/SPI-11 GFP-
G
SPI-2GFP-
SPI-2 GFP+
SPI-2 GFP+/SPI-2GFP-
H
SPI-12+oaf GFP-
SPI-12+oaf GFP+
SPI-12+oafGFP+/SPI-12+oaf GFP-

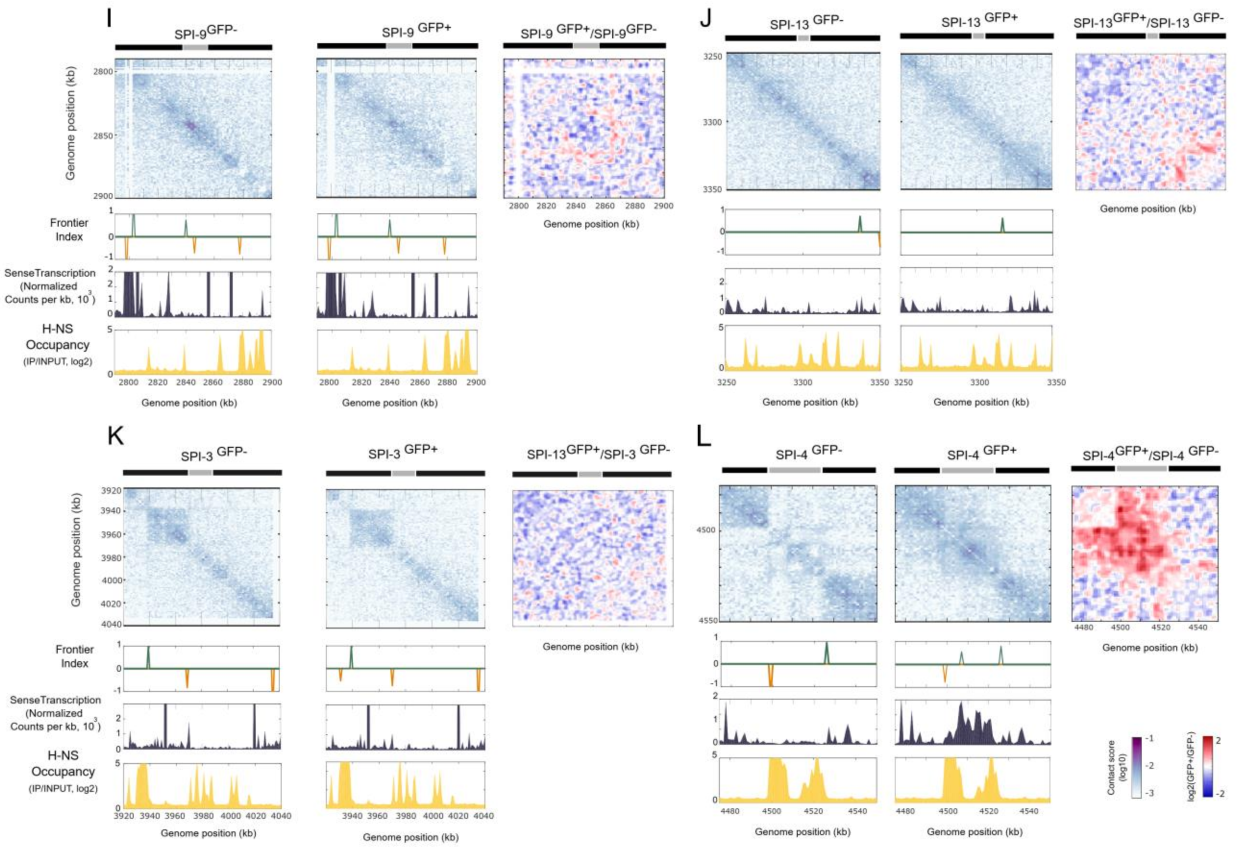


**Supplementary Figure 2: Chromatin organization and gene expression in other SPIs.**

A) Ratio of normalized contact maps for $GFP^+$ and $GFP^-$ populations (log2). A decrease or increase in contacts in the $GFP^+$ population compared with the $GFP^-$ population is represented with a blue or red color, respectively. White indicates no differences between the two populations.

B) to L) Contact maps, Frontier Index, gene expression and H-NS occupancy in the $GFP^-$ (left panel), $GFP^+$ (right panel), and normalized contact ratios centered in the indicated SPIs. Please note that the $P_{prgH}$-*gfp* reporter cassette used to sort $GFP^+$ cells is located downstream SPI-5, in between the region in which SPI-5 forms a loop.

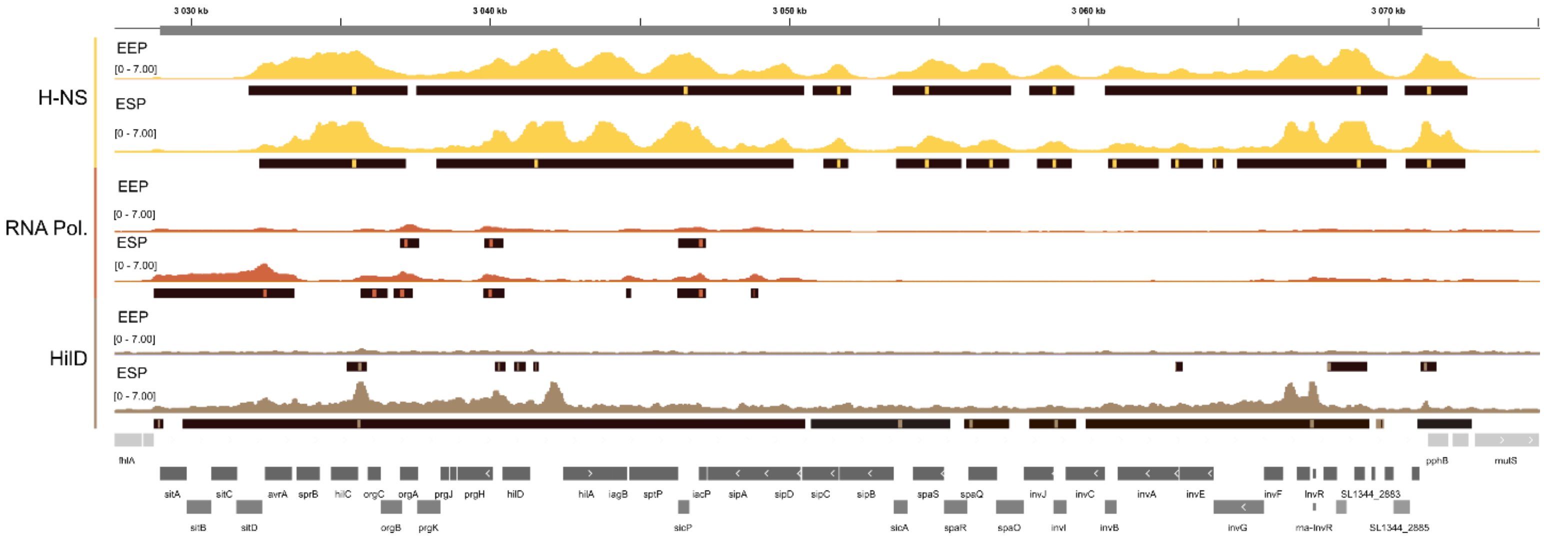


**Supplementary Figure 3: Protein occupancy within SPI-1 in non-sorted *Salmonella* populations**.

IGV snapshots centered on SPI-1 showing the binding occupancy of H-NS, RNA Polymerase (RNA Pol.) and HilD in repressed (EEP) or active SPI-1 (ESP) conditions. The histograms show the log2 (ChIP/Input) for each binding factor. Below the histograms, significant peaks detected by MACS2 are highlighted in black. Bottom plot shows SPI-1 genes and their neighboring genetic context.

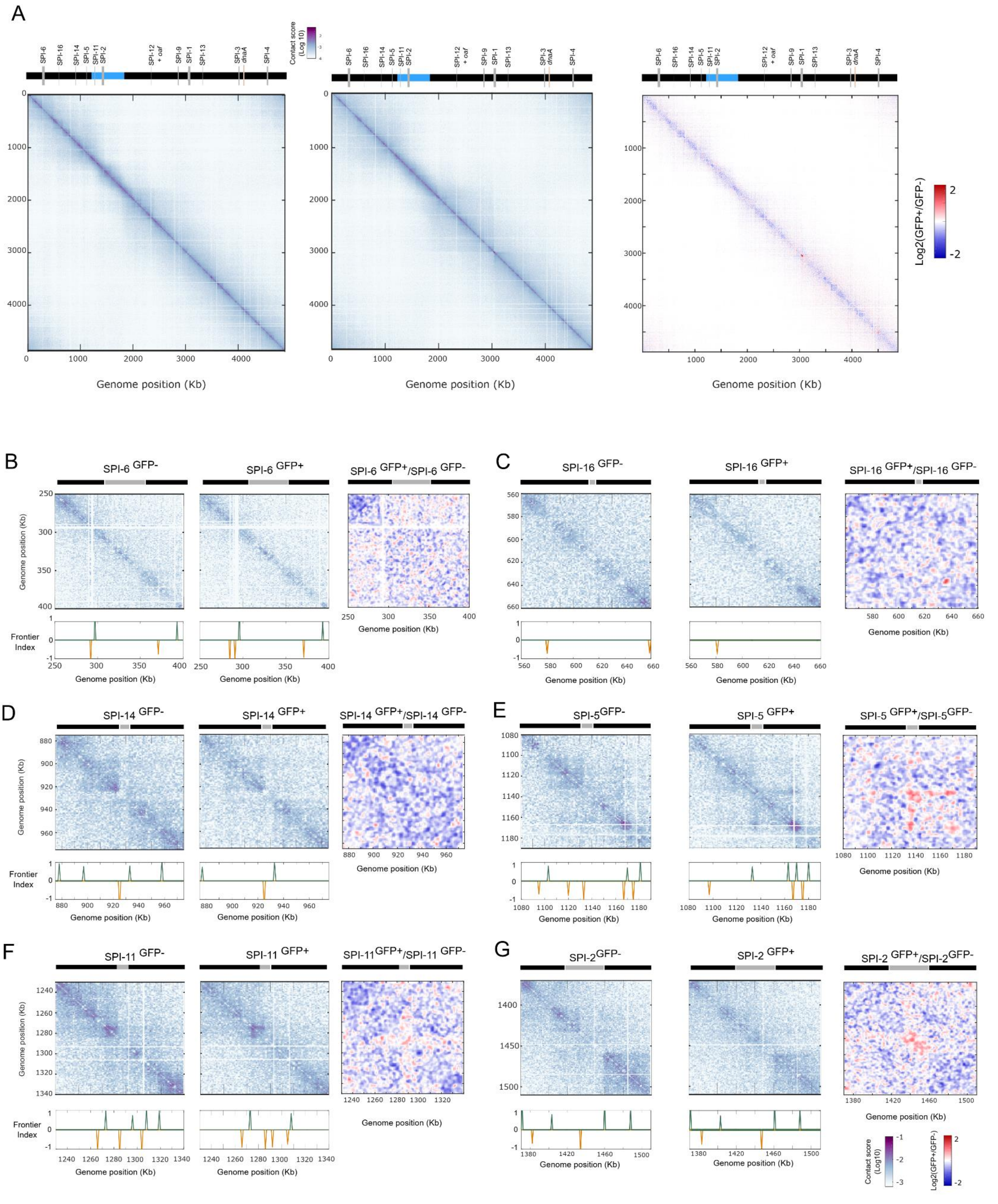

A
SPI-6
SPI-16
SPI-14
SPI-5
SPI-11
SPI-2
SPI-12 + oaf
SPI-9
SPI-1
SPI-13
SPI-3 dnaA
SPI-4
Contact score (Log 10)
Genome position (Kb)
Log2(GFP+/GFP-)
B
SPI-6 GFP-
SPI-6 GFP+
SPI-6 GFP+/SPI-6 GFP-
Frontier Index
C
SPI-16 GFP-
SPI-16 GFP+
SPI-16 GFP+/SPI-16 GFP-
D
SPI-14 GFP-
SPI-14 GFP+
SPI-14 GFP+/SPI-14 GFP-
E
SPI-5GFP-
SPI-5 GFP+
SPI-5 GFP+/SPI-5GFP-
F
SPI-11 GFP-
SPI-11 GFP+
SPI-11GFP+/SPI-11 GFP-
G
SPI-2GFP-
SPI-2 GFP+
SPI-2 GFP+/SPI-2GFP-
Genome position (Kb)

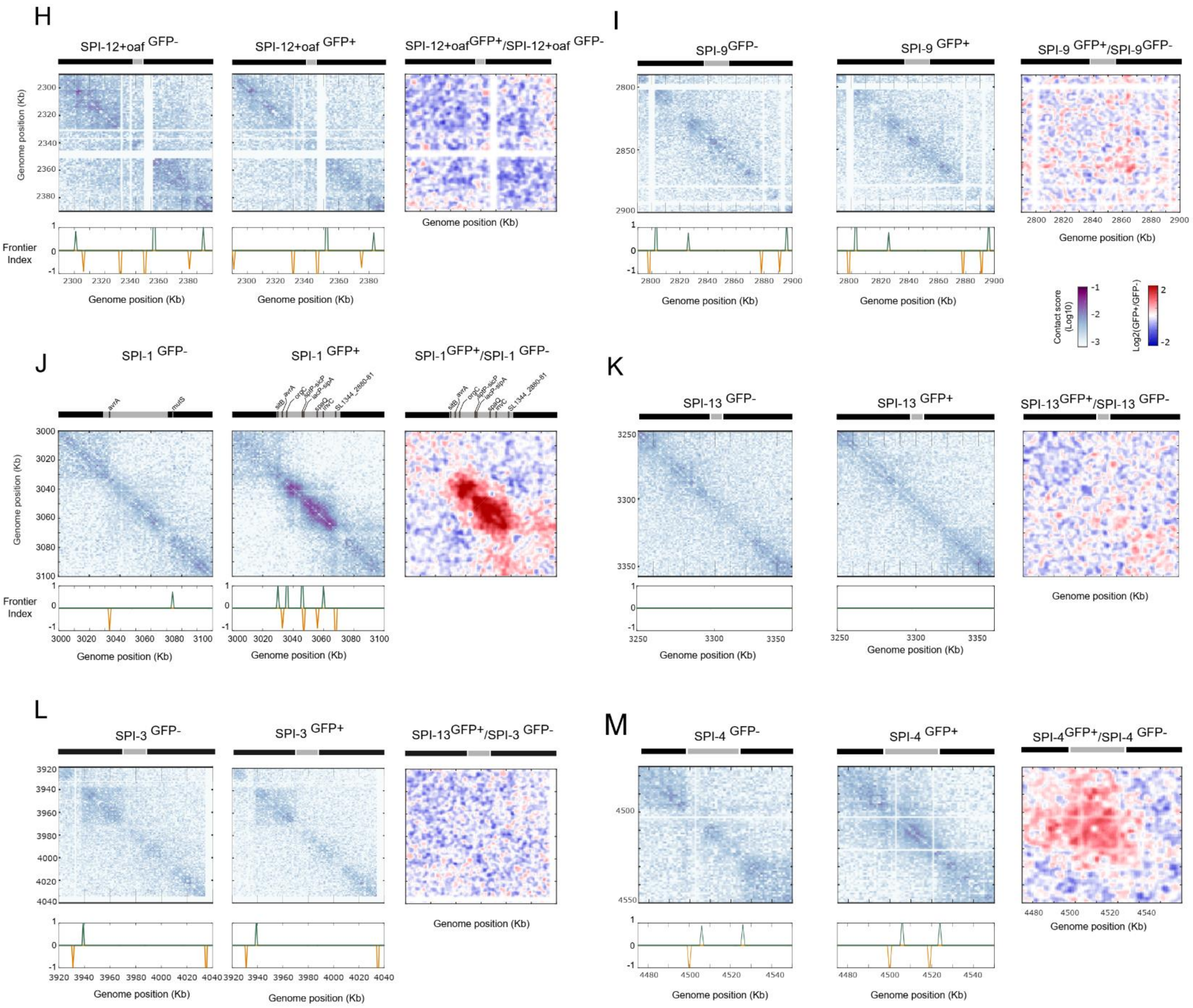


**Supplementary Figure 4. Chromosome organization in sorted GFP- and GFP+ populations in the absence of HilE**

A) 3D *Salmonella* chromosome folding in *hilE* deletion mutant cells in GFP- (Left panel) or GFP+ (middle panel) populations. Ratio of normalized contact maps for GFP+ and GFP- populations (Log2, right panel). A decrease or increase in contacts in the GFP+ population compared with the GFP-population is represented with a blue or red color, respectively. White indicates no differences between the two populations.

B) to M) Contact map, Frontier Index in the *hilE* GFP$^-$ (left panel), *hilE* GFP$^+$ population (middle panel), and normalized contact ratios (right panel) centered in all the SPIs.

Please note that the $P_{prgH}$-*gfp* reporter cassette used to sort $GFP^+$ cells is located downstream SPI-5, in between the region in which SPI-5 forms a loop.

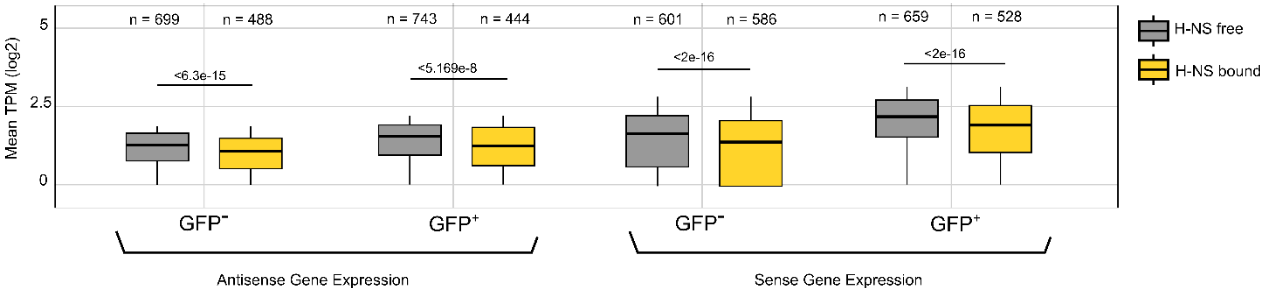


**Supplementary Figure 5: Level of transcription in low expression genes as a function of H-NS occupancy.** Silent or poorly expressed genes were identify as those which level of expression is in Category 1, which corresponds to the minimun of the boxplot of the mean transcripts per million (TPM, log 2) for the $GFP^-$ and $GFP^+$ populations in either sense or antisense direction (see Methods). In this figure, the level of gene expression is represented using boxplots, indicating the median (horizontal bar), the 25th and the 75th percentile (open box) and the rest of the population except for the outliers (whiskers). Values significantly different are indicated showing the *p*-value obtained for pairwise comparisons using the Wilcoxon rank sum test with continuity correction.

A

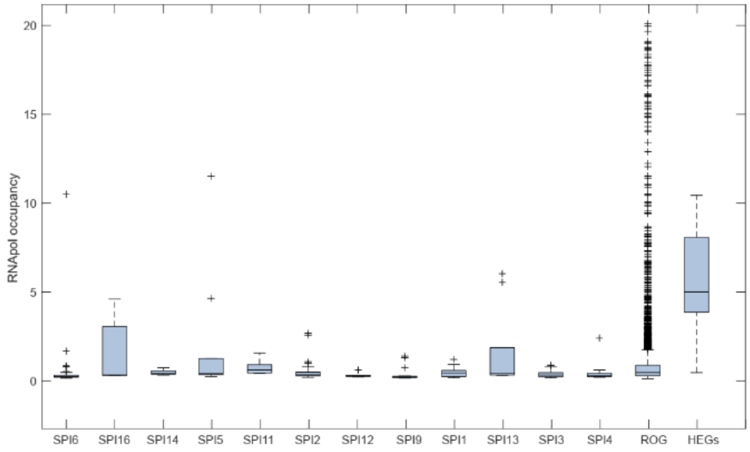


| | SPI6 | SPI16 | SPI14 | SPI5 | SPI11 | SPI2 | SPI12 | SPI9 | SPI1 | SPI13 | SPI3 | SPI4 | ROG |
|---|---|---|---|---|---|---|---|---|---|---|---|---|---|
| **ROG** | **2.9e-07** | 0.70984 | 0.93549 | 0.70984 | 0.19379 | 0.08060 | 0.14122 | **0.00070** | 0.11646 | 0.65410 | 0.06578 | **0.01369** | |
| **HEGs** | 2.3e-12 | 0.02106 | 7.4e-05 | 0.00548 | 0.00083 | 5.0e-12 | 7.3e-05 | 1.8e-08 | 7.1e-12 | 0.00466 | 1.2e-08 | 1.1e-11 | 1.0e-11 |

B

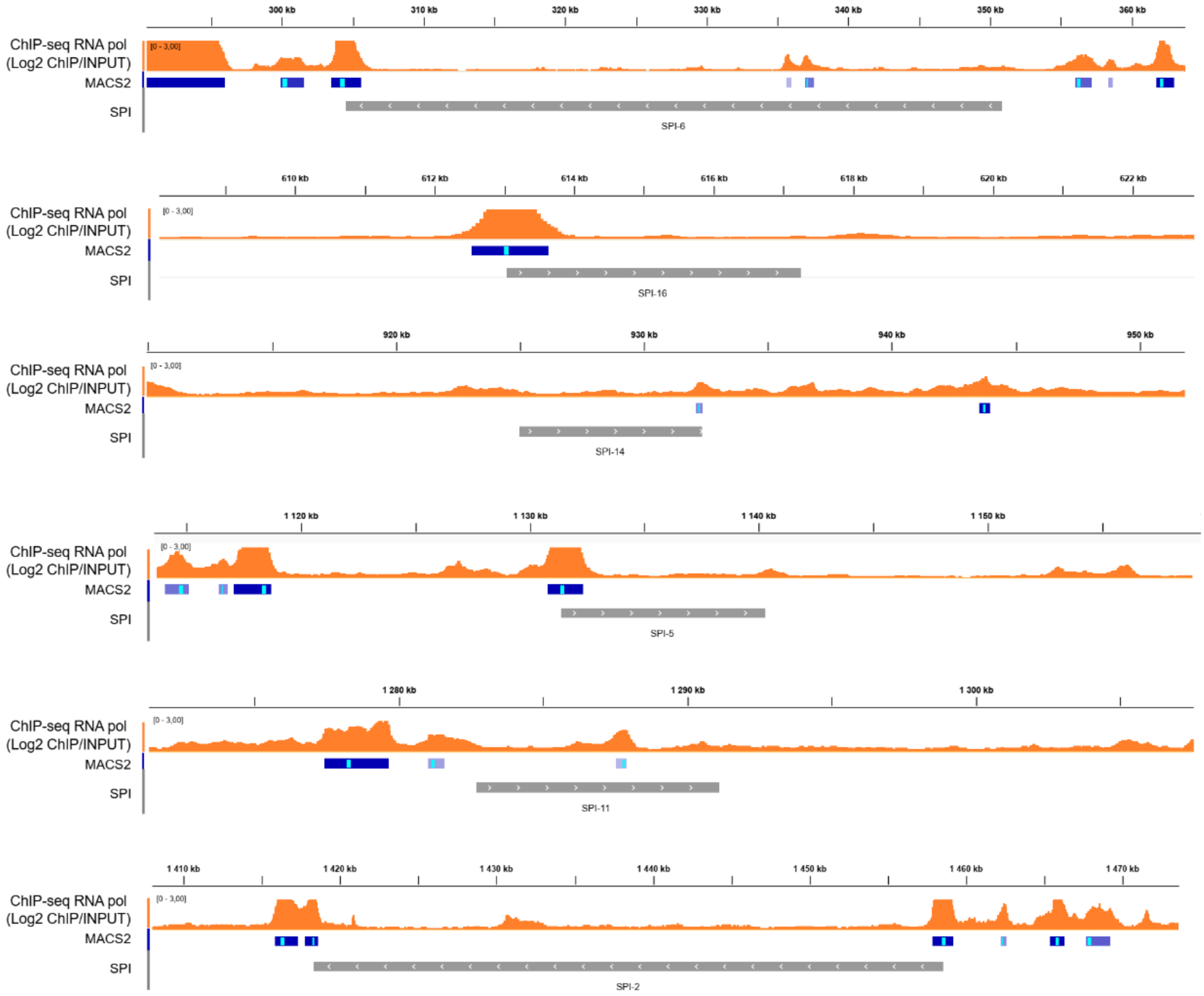

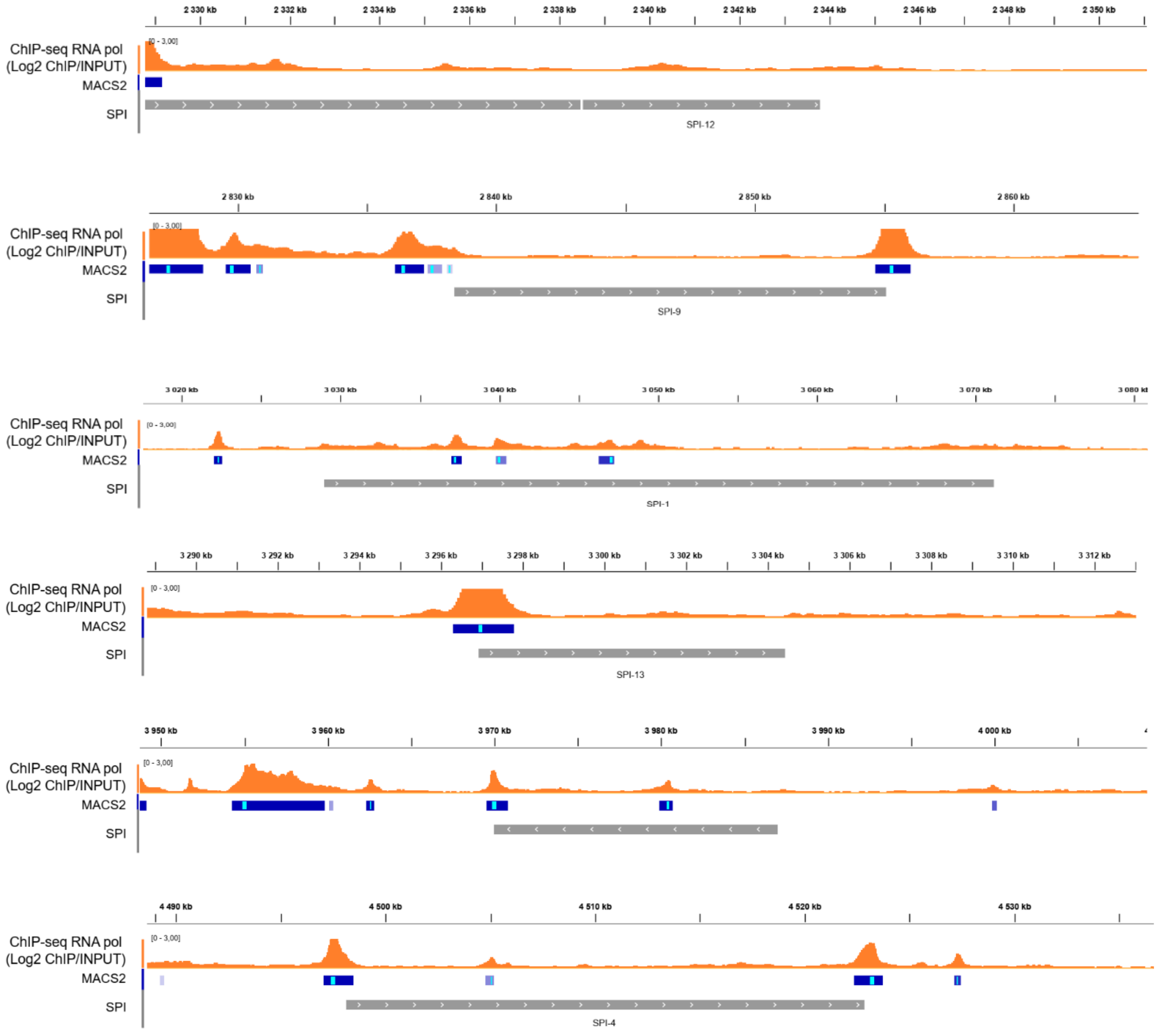


**Supplementary Figure 6. RNA polymerase occupancy near SPIs in exponentially growing *Salmonella* cells**.

A) Boxplot representing the RNA polymerase occupancy in silenced SPIs (EEP condition), the rest of genes of the chromosome (RoG) and highly expressed genes (HEGs). Bellow the boxplot, the Pairwise comparisons of the Wilcoxon rank sum exact test is shown. Significant values different than those of RoGs or HEGs are highlighted in bold in the table ($p$<0.05).

B) IGV snapshots centered on SPIs showing the binding occupancy of RNA polymerase in repressed (EEP) SPI-1 conditions.

A

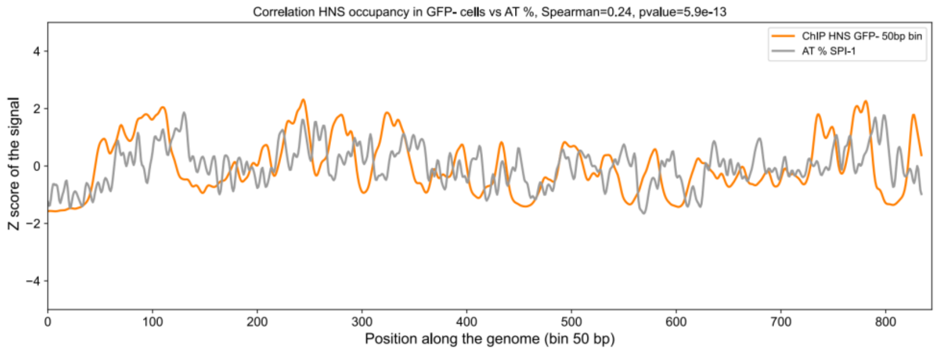


B

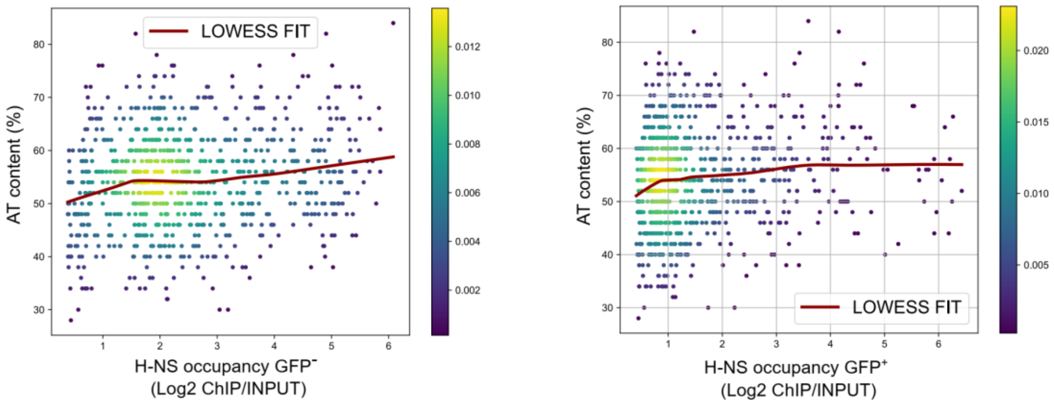


**Supplementary Figure 7: Correlation between H-NS occupancy and percentage of AT content within SPI-1 in sorted populations of *Salmonella*.**

A) Correlation between H-NS occupancy in the GFP$^-$ population (log2(ChIP/Input)) and AT content (%) within SPI-1. AT content was computed using a sliding window of 50 bp in size with an increasing step of 50bp.

B) H-NS abundance as a function of AT content in GFP$^-$ (left panel) or GFP$^+$ (Right panel) populations.

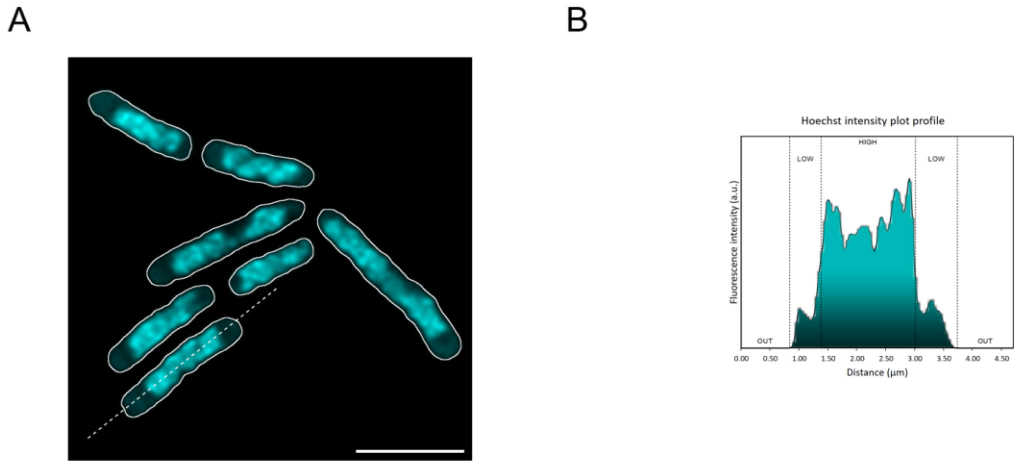


**Supplementary Figure 8. 3D SIM Imaging of *Salmonella*.**

A) Single plane SIM² image of Hoechst labelled bacterial DNA (scale bar = 2µm).
B) Related intensity plot profile, corresponding to the dotted line drawn on the image in (A).

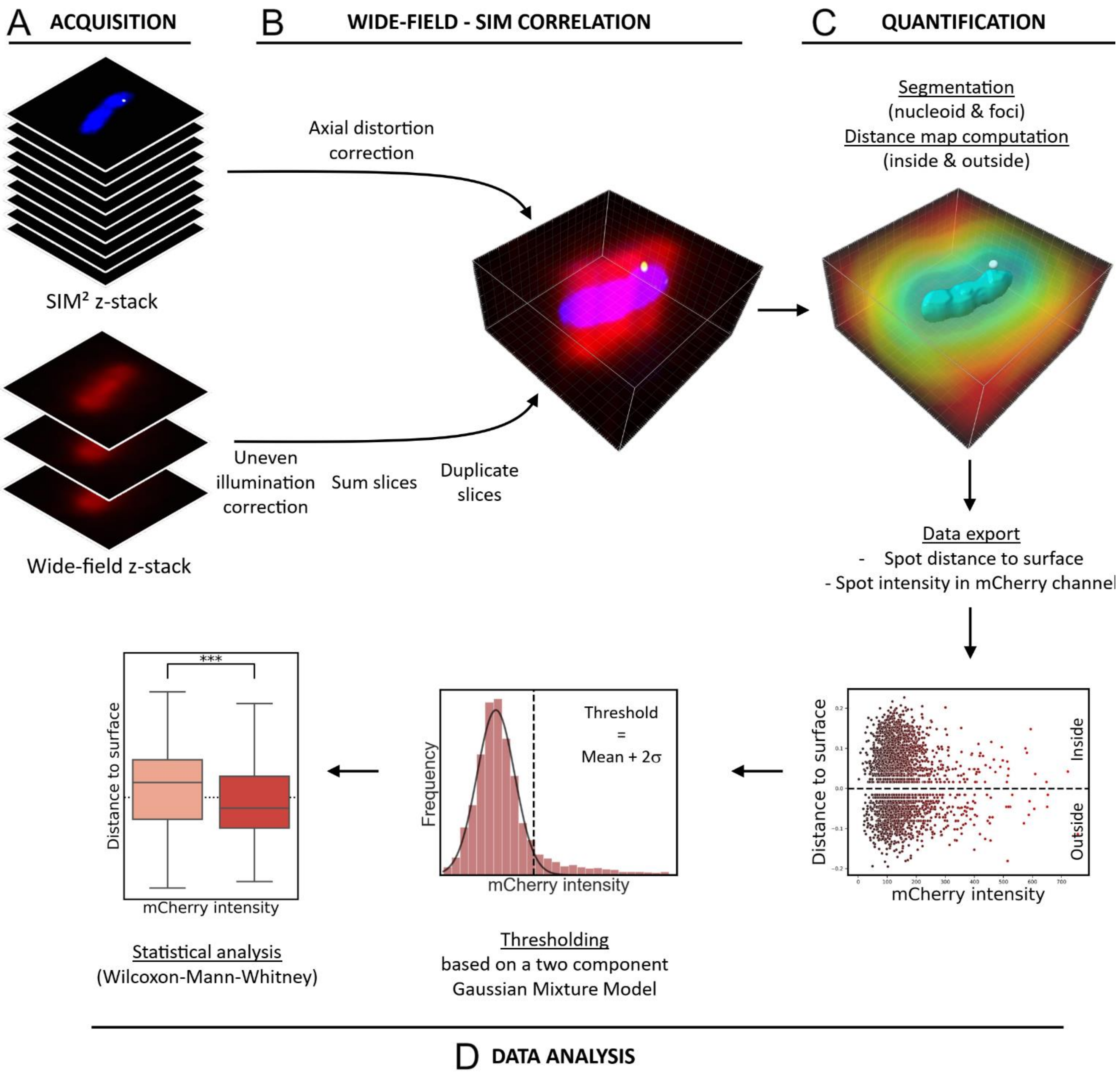


**Supplementary Figure 9. 3D SIM data analysis**.

A) Cells were acquired in Z stacks that included the whole bacteria, by switching between SIM$^2$ Z-stacks and wide-field Z-stacks.

B) Wide-Field & SIM correlation process: Following the acquisition steps, the wide-field and SIM$^2$ images were subjected to different processing procedures. The Hoechst and YFP SIM$^2$ image stacks were corrected for Z distortion. Z-axis intensity profiles were produced transversely to the bacteria (approximately 10 per field) in order to estimate the Z distortion after thresholding and fitting the resulting signals by an ellipse. This factor was used to correct the Z step and thus obtain isotropic cross-sections of the bacteria, limiting errors in 3D distance calculations. The mCherry wide-field stack, employed subsequently for the purpose of intensity quantification, was corrected for uneven illumination: a Rhodamine B solution was imaged under identical conditions to serve as a reference. After background subtraction, the images were divided by this reference image. To avoid errors in correlating the wide field data with the SIM data, a z-projection of the wide-field stack was performed and later duplicated multiple times in order to match the number of planes in the SIM image.

C) Quantification: Following the merging of the data, the data set was processed for segmentation. The Hoechst channel was treated to generate surfaces representing all regions of high DNA density, while the YFP channel was used to identify ParB:YFP/*parS*$^{pMT1}$ foci as spots. To remove aberrant foci detections, we kept them for the analysis only if they were closer than 0.2 µm from a nucleoid surface (representing more than 98% of the total detected foci). Two distance maps were then generated based on the surfaces previously generated, with measurements taken towards the inside and outside. The distance between a focus and the closest surface was obtained by measuring the value of each spot in the distance map channel. To determine whether a focus belonged to a SPI-1 ON or SPI-1 OFF cell, the value of the spot in the mCherry projection channel was measured.

D) Data Analysis: In order to identify the threshold between the SPI-1ON and SPI-1 OFF populations, a Gaussian Mixture Model based on two components (Python, Scikit-learn library) was employed on a histogram presenting spot intensities in the mCherry channel. The Gaussian of the lowest population (comprising 90% of the spots) was used to determine the threshold with precision by calculating the mean of the Gaussian fit plus twice its standard deviation. Following the splitting of the data into two populations, the spot distances from the DNA-dense regions were subjected to a statistical analysis using the Wilcoxon-Mann-Whitney test.

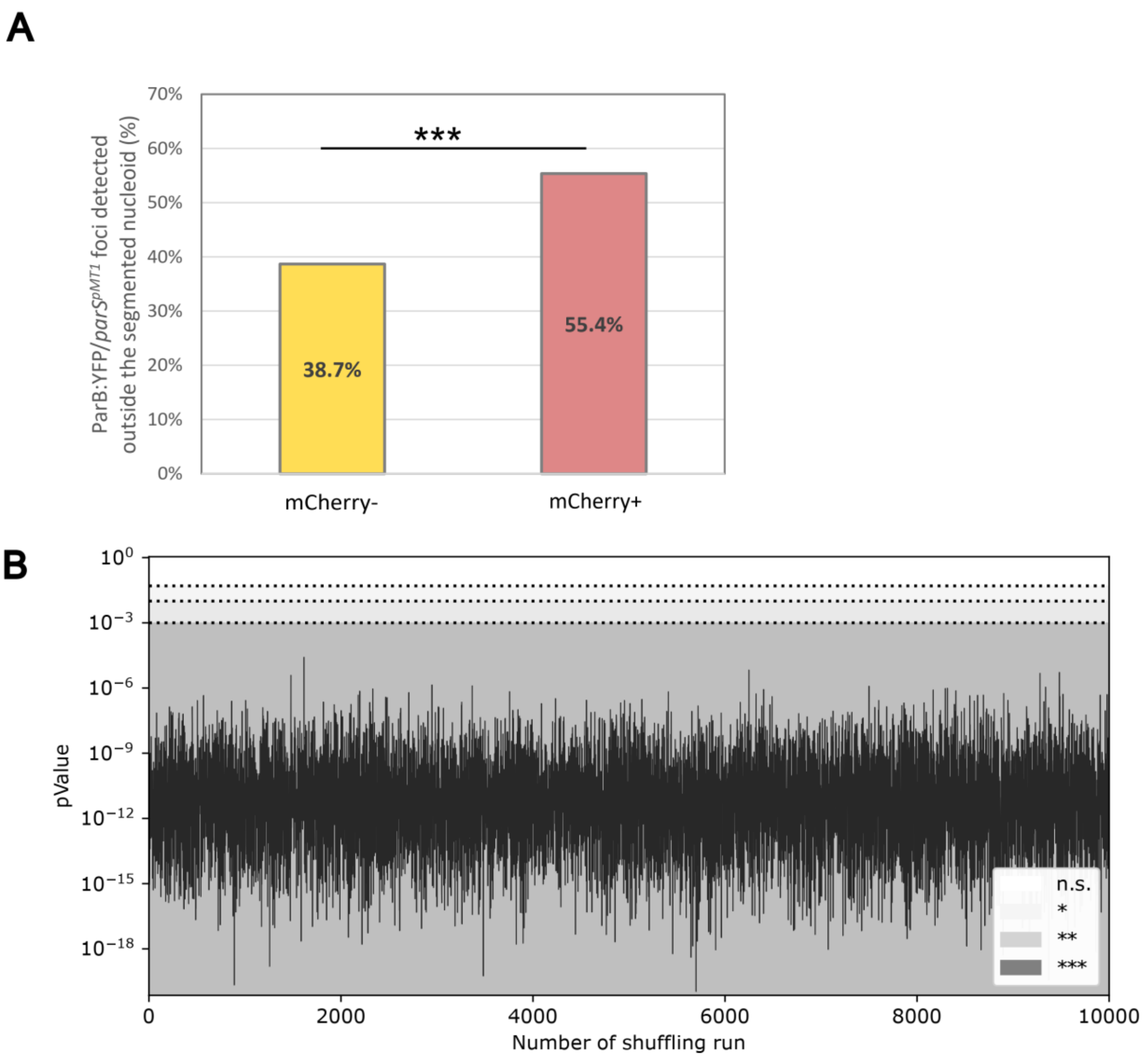


**Supplementary Figure 10: ParB:YFP/*parS*$^{pMT1}$ foci localization in *Salmonella* cells**

A) Histogram representing the fraction of the ParB:YFP/*parS*$^{pMT1}$ foci detected outside the segmented nucleoid region, in both mCherry$^-$ and mCherry$^+$ populations. Chi-square test showing the significance of the differences between the two populations (p-value = 2.485e-16).

B) Computation of the *p*-value, obtained from a Mann-Whitney test applied to the ParB:YFP/*parS*$^{pMT1}$ foci distances from the nucleoid surface in both mCherry$^-$ and mCherry$^+$. To avoid a side effect of the population size, we used the same number of foci for each population (n=652). To do so, the distances of all the mCherry$^+$ foci were compared to a random subpopulation of mCherry$^-$ foci (repeated 10,000 times). This approach demonstrates that disparities in population size do not influence the significance of the difference in localization between the two populations.

**Supplementary Table 1:** Percentage of $GFP^+$ cells in the different strains and conditions studied, as determined by FACS

| **Strain** | **$GFP^+$ in ESP Replicate 1** | **$GFP^+$ in ESP Replicate 2** |
|---|---|---|
| SL1344 $P_{prgH}$-*gfp* | 11.03% | 9.5% |
| SL1344 *hilD* $P_{prgH}$-*gfp* | 0.31% | 0.51% |
| SL1344 *hilE* $P_{prgH}$-*gfp* | 27.77% | 25.92% |

**Supplementary Table 2:** Main functions of SPIs encoded in S. Typhimurium SL1344

| ***Salmonella* Pathogenicity Island** | **Function** | **Co-regulated with SPI-1** | **Reference** |
|---|---|---|---|
| SPI-6 | Encodes a type VI secretion system (T6SS) that enables gut colonization by killing commensal bacteria, it also contributes to intracellular replication and systemic dissemination in mice. | NR | [3,4] |
| SPI-16 | Encodes genes responsible for O-antigen glucosylation, enhancing long-term intestinal persistence and fecal shedding of *Salmonella* in mice | NR | [5] |
| SPI-14 | Encodes *loiA*, which regulates SPI-1 activation in low oxygen conditions, promoting *Salmonella* intestinal invasion. | Yes | [6] |
| SPI-5 | Encodes effectors involved in different stages of infection: SopB, secreted by T3SS-1, promotes invasion, while PipB, translocated by T3SS-2, promotes intra-macrophage survival. | Yes | [7] |
| SPI-11 | Encodes several genes that are critical for *Salmonella* survival within macrophages**.** | NR | [8,9] |
| SPI-2 | Encodes for Type 3 secretion system (T3SS-2) and effectors essential for survival in macrophages. | Yes | [10] |
| SPI-12 | Encodes *sspH2*, secreted by T3SS-2, which contributes to immune evasion and intracellular survival. | NR | [11] |
| SPI-9 | SPI-9 in *S.* Typhimurium is not well characterized, but in *S. Typhi*, it encodes a Type 1 secretion system and an adhesin protein involved in epithelial cell adhesion under high osmolarity and low pH conditions. | NR | [12] |
| SPI-1 | Encodes for Type 3 secretion system (T3SS-1) and effectors essential for invasion and cytosolic lifestyle. | NA | [13] |
| SPI-13 | Encodes genes essential for virulence in mice and survival within macrophages. | NR | [14,15] |
| SPI-3 | Encodes the *mgtCB* gene, a high-affinity Mg2+ uptake system required for *Salmonella* survival in macrophages and low Mg2+ environments. It also contains an adhesin essential for fibronectin binding and intestinal persistence. | NR | [16,17] |

| | | | |
|---|---|---|---|
| SPI-4 | Encodes a type 1 secretion system and a major adhesin protein SiiE, which are important for bacterial entry into epithelial cells by promoting clustering and membrane ruffling at the invasion site. | Yes | [18,19] |

NR: Not Reported in *S.* Typhimurium
NA: Not Applicable

**Supplementary Table 3:** SPI encoded genes upregulated in the GFP+ population

| SPI | Gene name | Encoded Product | KEGG Orthology (KO) | KO sub category |
|---|---|---|---|---|
| SPI-5 | *pipC* | Cell invasion protein | not assigned | not assigned |
| SPI-5 | *sopB* | Type III secretion system effector protein | Human Diseases | Infectious disease: bacterial |
| SPI-11 | *SL1344_1177* | Predicted bacteriophage protein | not assigned | not assigned |
| SPI-11 | *pagC* | Outer membrane invasion protein (PagC) | Environmental Information Processing | Signal transduction |
| SPI-2 | *ssrB* | Two-component system, LuxR family, secretion system response regulator SsrB | Environmental Information Processing | Signal transduction |
| SPI-1 | *avrA* | Type III secretion system (T3SS) effector protein-regulator of *Salmonella*-induced inflammatory response | Human Diseases | Infectious disease: bacterial |
| SPI-1 | *sprB* | AraC-family transcriptional regulator | not assigned | not assigned |
| SPI-1 | *hilC* | AraC-family transcriptional regulator | not assigned | not assigned |
| SPI-1 | *orgC* | Type III secretion system effector protein | not assigned | not assigned |
| SPI-1 | *orgB* | Component of the sorting Platform of the T3SS1 | not assigned | not assigned |
| SPI-1 | *orgA* | Component of the sorting Platform of the T3SS1 | not assigned | not assigned |
| SPI-1 | *prgK* | T3SS1 inner membrane ring protein | Brite Hierarchies | Protein families: signaling and cellular processes |
| SPI-1 | *prgJ* | Component of the T3SS1 | Human Diseases | Infectious disease: bacterial |
| SPI-1 | *prgH* | Component of the T3SS1 | Brite Hierarchies | Protein families: signaling and cellular processes |

| | | | | |
|---|---|---|---|---|
| SPI-1 | *hilD* | AraC-family transcriptional regulator. SPI-1 master regulator | not assigned | not assigned |
| SPI-1 | *hilA* | Transcriptional regulator HilA, main transcriptional regulator of SPI1 | Brite Hierarchies | Protein families: genetic information processing |
| SPI-1 | *sptP* | T3SS1 effector protein | Human Diseases | Infectious disease: bacterial |
| SPI-1 | *sicP* | Chaperone (associated with virulence) | Not assigned | not assigned |
| SPI-1 | *iacP* | Required for the invasion of nonphagocytic cells. | Metabolism | Lipid metabolism |
| SPI-1 | *sipA* | T3SS1 effector protein-involved in actin bundling and polymerisation leading to epithelial cell invasion and formation of the SCV | Human Diseases | Infectious disease: bacterial |
| SPI-1 | *sipD* | Part of the T3SS1 Translocon | Human Diseases | Infectious disease: bacterial |
| SPI-1 | *sipC* | T3SS1 effector protein-involved in bacterial entry by actin bundling and part of the Translocon | Human Diseases | Infectious disease: bacterial |
| SPI-1 | *sipB* | T3SS1 effector protein | Human Diseases | Infectious disease: bacterial |
| SPI-1 | *sicA* | T3SS1 -associated chaperone | not assigned | not assigned |
| SPI-1 | *spaS* | T3SS1 export apparatus switch protein | Brite Hierarchies | Protein families: signaling and cellular processes |
| SPI-1 | *spaR* | T3SS1 export apparatus protein | Brite Hierarchies | Protein families: signaling and cellular processes |
| SPI-1 | *spaQ* | T3SS1 export apparatus protein | Brite Hierarchies | Protein families: signaling and cellular processes |
| SPI-1 | *spaP* | T3SS1 export apparatus protein | Brite Hierarchies | Protein families: signaling and cellular processes |
| SPI-1 | *spaO* | Component of the sorting Platform of the T3SS1 | Environmental Information Processing | Membrane transport |

| | | | | |
|---|---|---|---|---|
| SPI-1 | *invJ* | T3SS1 protein | Brite Hierarchies | Protein families: signaling and cellular processes |
| SPI-1 | *invC* | component of the T3SS1 sorting platform | Brite Hierarchies | Protein families: signaling and cellular processes |
| SPI-1 | *invB* | T3SS1 chaperone | Brite Hierarchies | Protein families: signaling and cellular processes |
| SPI-1 | *invE* | T3SS1 protein | Brite Hierarchies | Protein families: signaling and cellular processes |
| SPI-1 | *invG* | T3SS1 outer membrane ring protein | Brite Hierarchies | Protein families: signaling and cellular processes |
| SPI-1 | *InvR* | small RNA | not assigned | not assigned |
| SPI-3 | *cigR* | hypothetical inner membrane protein | not assigned | not assigned |
| SPI-4 | *siiB* | inner membrane protein | not assigned | not assigned |
| SPI-4 | *siiC* | Component of a type-I secretion system 4 (T1SS4) | Brite Hierarchies | Protein families: signaling and cellular processes |
| SPI-4 | *siiD* | Component of T1SS4 | Brite Hierarchies | Protein families: signaling and cellular processes |

**Supplementary Table 4**. Strains used in this study

| Name | Description | Source or Reference |
|---|---|---|
| Wild-Type | *Salmonella enterica* serovar Typhimurium SL1344 | [20] |
| MKB1 | $P_{prgH}$-*gfp* $Cm^R$ | [21] |
| MKB7 | *hilE*::$cm^R$ | This study |
| MKB13 | *hns-3xFLAG* $Cm^R$ | This study |
| MKB15 | *hilE* $P_{prgH}$-*gfp* $Cm^R$ | This study |
| MKB16 | *hilD*::$Cm^R$ | This study |
| MKB17 | *rpoC-3xFLAG* $Cm^R$ | This study |
| MKB18 | *hilD-3xFLAG* $Cm^R$ | This study |
| MKB27 | *hns-3xFLAG* $P_{prgH}$-*gfp* $Cm^R$ | This study |
| MKB33 | *prgH(stop)mCherry* $Kn^R$ $parS^{pMT1}Cm^R$ | This study |

**Supplementary Table 5**. Plasmids used in this study

| Name | Description | Source or Reference |
|---|---|---|
| pKD3 | PCR template plasmid containing a chloramphenicol resistance cassette. | [22] |
| pKD4 | PCR template plasmid containing a kanamycin resistance cassette. | [22] |
| pKD46 | Plasmid encoding λ-Red system for recombination. | [22] |
| pCP20 | Plasmid encoding FLP recombinase. | [22] |
| pMKB1 | PCR template plasmid containing promoter less *mCherry.* | This study |
| pSPB5 | $P_{lac}$ $parB^{pMT1}$:*yfp* $amp^R$ | This study |

**Supplementary Table 6.** Hi-C libraries performed in this study

| Id | Library | Conditions | Total pairs processed | Valid interactions |
|---|---|---|---|---|
| MKL3 | WT EEP rep1 | growth in LB at 37°C EEP (OD 0.1) | 6273869 | 5359451 |
| MKL8 | WT EEP rep2 | | 9180949 | 8922179 |
| MKL19 | WT EEP rep3 | | 10830147 | 6330485 |
| WT EEP merged | **WT EEP merge** | | 26336107 | **20 649 013** |
| MKL1 | WT ESP rep1 | growth in LB at 37°C ESP (OD 2) | 7837734 | 7154256 |
| MKL4 | WT ESP rep2 | | 6893876 | 6688347 |
| MKL9 | WT ESP rep3 | | 8973183 | 5648927 |
| MKL20 | WT ESP rep3 | | 11064794 | 9626700 |
| WT ESP merged | **WT ESP merge** | | 34663755 | **29 024 094** |
| MKL15 | WT Anaerobic growth | Anaerobic growth in LB at 37°C to OD 0.3 | 4077901 | 3 999 140 |
| MKL15_bis* | | | 1299562 | 1 292 701 |
| MKL15_Merged | **WT Anaerobic growth merged** | | 5379716 | **5 263 346** |
| MKL16 | WT Oxygen shock | Static growth in LB at 37°C to OD 0.3 then 15 min aerobic growth | 2802296 | 2 754 855 |
| MKL16_bis* | | | 3544955 | 3 515 850 |
| MKL16_Merged | **WT Oxygen shock merged** | | 6344054 | **6 232 640** |
| MKL41 | **SL1344 *hilE* $P_{prgH}$-*gfp* ESP GFP+** | growth in LB at 37°C ESP (OD 2) Sorted only GFP positive population | 6555694 | **5 815 621** |
| MKL42 | **SL1344 *hilE* $P_{prgH}$-*gfp* ESP GFP-** | growth in LB at 37°C ESP (OD 2) Sorted only GFP negative population | 5921230 | **5 622 714** |
| MKL72 | SL1344 *hilD* ESP rep1 | growth in LB at 37°C ESP (OD 2) | 7282238 | 6 956 139 |
| MKL134 | SL1344 *hilD* ESP rep2 | | 6191117 | 5 883 056 |
| DhilD_ESP_merged | **SL1344 *hilD* ESP merged** | | 13475614 | **12 839 983** |
| MKL81 | SL1344 $P_{prgH}$-*gfp* ESP GFP+ | growth in LB at 37°C ESP (OD 2) Sorted only GFP positive population | 2271159 | 2 135 781 |
| MKL81_bis* | | | 10432522 | 8 350 519 |
| MKL81_Merged | **SL1344 $P_{prgH}$-*gfp* ESP GFP+ Merged** | | 12703915 | **9 758 492** |
| MKL82 | SL1344 $P_{prgH}$-*gfp* ESP GFP+ | growth in LB at 37°C ESP (OD 2) Sorted only GFP negative population | 1973214 | 1 920 635 |
| MKL82_bis* | | | 9032070 | 8 403 073 |
| MKL82_Merged | **SL1344 $P_{prgH}$-*gfp* ESP GFP+** | | 11005464 | **10 109 980** |

*Libraries were re-sequenced to have higher valid reads number.

**Supplementary Table 7.** ChIP-seq libraries performed in this study

| ID | Library | Conditions | Mapped_reads_count |
|---|---|---|---|
| MKL91 | Input SL1344 *hilD*-3xFLAG EEP rep1 | growth in LB at 37°C EEP (OD 0.1) | 36194599 |
| MKL93 | Input SL1344 *hilD*-3xFLAG EEP rep2 | | 31646346 |
| **merge** | **Input SL1344 *hilD*-3xFLAG EEP merge** | | **67 840 945** |
| MKL92 | Input SL1344 *hilD*-3xFLAG ESP rep1 | growth in LB at 37°C ESP (OD 2) | 36937073 |
| MKL94 | Input SL1344 *hilD*-3xFLAG ESP rep2 | | 29872125 |
| merge | **Input SL1344 *hilD*-3xFLAG ESP merge** | | **66 809 198** |
| MKL95 | IP SL1344 *hilD*-3xFLAG EEP rep1 | growth in LB at 37°C EEP (OD 0.1) | 11138115 |
| MKL97 | IP SL1344 *hilD*-3xFLAG EEP rep2 | | 10560086 |
| merge | **IP SL1344 *hilD*-3xFLAG EEP merge** | | **21 698 201** |
| MKL96 | IP SL1344 *hilD*-3xFLAG ESP rep1 | growth in LB at 37°C ESP (OD 2) | 14354457 |
| MKL98 | IP SL1344 *hilD*-3xFLAG ESP rep2 | | 9670734 |
| | **IP SL1344 *hilD*-3xFLAG ESP merge** | | **24 025 191** |
| MKL99 | Input SL1344 *rpoC*-3xFLAG EEP rep1 | growth in LB at 37°C EEP (OD 0.1) | 29246161 |
| MKL101 | Input SL1344 *rpoC*-3xFLAG EEP rep2 | | 26394520 |
| MKL135 | Input SL1344 *rpoC*-3xFLAG EEP rep3 | | 20486978 |
| MKL137 | Input SL1344 *rpoC*-3xFLAG EEP rep4 | | 23789930 |
| **merge** | **Input SL1344 *rpoC*-3xFLAG EEP merge** | | **99 917 589** |
| MKL103 | IP SL1344 *rpoC*-3xFLAG EEP rep1 | growth in LB at 37°C EEP (OD 0.1) | 20969598 |
| MKL105 | IP SL1344 *rpoC*-3xFLAG EEP rep2 | | 15991264 |
| MKL139 | IP SL1344 *rpoC*-3xFLAG EEP rep3 | | 15894320 |
| MKL141 | IP SL1344 *rpoC*-3xFLAG EEP rep4 | | 16582489 |
| **merge** | **IP SL1344 *rpoC*-3xFLAG EEP merge** | | **69 437 671** |
| MKL107 | Input SL1344 *hns*-3xFLAG EEP rep1 | growth in LB at 37°C EEP (OD 0.1) | 31922981 |
| MKL109 | Input SL1344 *hns*-3xFLAG EEP rep2 | | 27133135 |
| **merge** | **Input SL1344 *hns*-3xFLAG EEP merge** | | **59 056 116** |
| MKL108 | Input SL1344 *hns*-3xFLAG ESP rep1 | growth in LB at 37°C ESP (OD 2) | 20217682 |

| | | | |
|---|---|---|---|
| MKL110 | Input SL1344 *hns*-3xFLAG ESP rep2 | | 25978731 |
| **merge** | **Input SL1344 *hns*-3xFLAG ESP merge** | | **46 196 413** |
| MKL111 | IP SL1344 hns-3xFLAG EEP rep1 | growth in LB at 37°C EEP (OD 0.1) | 25621825 |
| MKL113 | IP SL1344 hns-3xFLAG EEP rep2 | | 28369797 |
| **merge** | **IP SL1344 hns-3xFLAG EEP merge** | | **53 991 622** |
| MKL112 | IP SL1344 hns-3xFLAG ESP rep1 | growth in LB at 37°C ESP (OD 2) | 18856253 |
| MKL114 | IP SL1344 hns-3xFLAG ESP rep2 | | 30330205 |
| **merge** | **IP SL1344 hns-3xFLAG ESP merge** | | **49 186 458** |
| MKL115 | Input SL1344 hns-3xFLAG $P_{prgH}$-*gfp* ESP GFP+ rep1 | growth in LB at 37°C ESP (OD 2) Sorted only GFP positive population | 33930348 |
| MKL117 | Input SL1344 hns-3xFLAG $P_{prgH}$-*gfp* ESP GFP+ rep2 | | 29563335 |
| **merge** | **Input SL1344 hns-3xFLAG $P_{prgH}$-*gfp* ESP GFP+ merge** | | **63 493 683** |
| MKL116 | Input SL1344 hns-3xFLAG $P_{prgH}$-*gfp* ESP GFP- rep1 | growth in LB at 37°C ESP (OD 2) Sorted only GFP negative population | 38986395 |
| MKL118 | Input SL1344 hns-3xFLAG $P_{prgH}$-*gfp* ESP GFP- rep2 | | 36089550 |
| **merge** | **Input SL1344 hns-3xFLAG $P_{prgH}$-*gfp* ESP GFP- merge** | | **75 075 945** |
| MKL119 | IP SL1344 hns-3xFLAG $P_{prgH}$-*gfp* ESP GFP+ rep1 | growth in LB at 37°C ESP (OD 2) Sorted only GFP positive population | 56103616 |
| MKL121 | IP SL1344 hns-3xFLAG $P_{prgH}$-*gfp* ESP GFP+ rep2 | | 32316035 |
| **merge** | **IP SL1344 hns-3xFLAG $P_{prgH}$-*gfp* ESP GFP+ merge** | | **88 419 651** |
| MKL120 | IP SL1344 hns-3xFLAG $P_{prgH}$-*gfp* ESP GFP- rep1 | growth in LB at 37°C ESP (OD 2) Sorted only GFP negative population | 45910688 |
| MKL122 | IP SL1344 hns-3xFLAG $P_{prgH}$-*gfp* ESP GFP- rep2 | | 67132426 |
| **merge** | **IP SL1344 hns-3xFLAG $P_{prgH}$-*gfp* ESP GFP- merge** | | **113 043 114** |

**Supplementary Table 8.** RNA-seq libraries performed in this study

| ID | Library | Conditions | Total reads | Total mapped reads |
|---|---|---|---|---|
| MKR21 | Total RNA, SL1344 $P_{prgH}$-*gfp* rep1 | growth in LB at 37°C ESP (OD 2), sorted, $GFP^{+}$ population | 34 852 819 | 13 648 886 |
| MKR23 | Total RNA, SL1344 $P_{prgH}$-*gfp* rep2 | | 43 388 497 | 19 678 566 |
| MKR22 | Total RNA, SL1344 $P_{prgH}$-*gfp* rep1 | growth in LB at 37°C ESP (OD 2),sorted, $GFP^{-}$ population | 37 839 496 | 12 973 695 |
| MKR24 | Total RNA, SL1344 $P_{prgH}$-*gfp* rep2 | | 28 128 599 | 11 989 278 |

### References


1. Kröger, C. *et al.* An infection-relevant transcriptomic compendium for Salmonella enterica Serovar Typhimurium. *Cell Host Microbe* **14**, 683–695 (2013).
2. Lioy, V. S., Junier, I., Lagage, V., Vallet, I. & Boccard, F. Distinct Activities of Bacterial Condensins for Chromosome Management in Pseudomonas aeruginosa. *Cell Rep.* **33**, 108344 (2020).
3. Mulder, D. T., Cooper, C. A. & Coombes, B. K. Type VI secretion system-associated gene clusters contribute to pathogenesis of Salmonella enterica serovar Typhimurium. *Infect. Immun.* **80**, 1996–2007 (2012).
4. Sana, T. G. *et al.* Salmonella Typhimurium utilizes a T6SS-mediated antibacterial weapon to establish in the host gut. *Proc. Natl. Acad. Sci. U. S. A.* **113**, E5044-5051 (2016).
5. Bogomolnaya, L. M., Santiviago, C. A., Yang, H.-J., Baumler, A. J. & Andrews-Polymenis, H. L. 'Form variation' of the O12 antigen is critical for persistence of Salmonella Typhimurium in the murine intestine. *Mol. Microbiol.* **70**, 1105–1119 (2008).
6. Jiang, L. *et al.* Signal transduction pathway mediated by the novel regulator LoiA for low oxygen tension induced Salmonella Typhimurium invasion. *PLoS Pathog.* **13**, e1006429 (2017).
7. Knodler, L. A. *et al.* Salmonella effectors within a single pathogenicity island are differentially expressed and translocated by separate type III secretion systems. *Mol. Microbiol.* **43**, 1089–1103 (2002).

8. Miller, S. I., Kukral, A. M. & Mekalanos, J. J. A two-component regulatory system (phoP phoQ) controls Salmonella typhimurium virulence. *Proc. Natl. Acad. Sci. U. S. A.* **86**, 5054–5058 (1989).

9. Lee, Y. H., Kim, S., Helmann, J. D., Kim, B.-H. & Park, Y. K. RaoN, a small RNA encoded within Salmonella pathogenicity island-11, confers resistance to macrophage-induced stress. *Microbiol. Read. Engl.* **159**, 1366–1378 (2013).

10. Hensel, M. *et al.* Genes encoding putative effector proteins of the type III secretion system of Salmonella pathogenicity island 2 are required for bacterial virulence and proliferation in macrophages. *Mol. Microbiol.* **30**, 163–174 (1998).

11. McGhie, E. J., Brawn, L. C., Hume, P. J., Humphreys, D. & Koronakis, V. Salmonella takes control: effector-driven manipulation of the host. *Curr. Opin. Microbiol.* **12**, 117–124 (2009).

12. Velásquez, J. C. *et al.* SPI-9 of Salmonella enterica serovar Typhi is constituted by an operon positively regulated by RpoS and contributes to adherence to epithelial cells in culture. *Microbiology* **162**, 1367–1378 (2016).

13. Raffatellu, M. *et al.* SipA, SopA, SopB, SopD, and SopE2 contribute to Salmonella enterica serotype typhimurium invasion of epithelial cells. *Infect. Immun.* **73**, 146–154 (2005).

14. Shi, L. *et al.* Proteomic analysis of Salmonella enterica serovar typhimurium isolated from RAW 264.7 macrophages: identification of a novel protein that contributes to the replication of serovar typhimurium inside macrophages. *J. Biol. Chem.* **281**, 29131–29140 (2006).

15. Haneda, T., Ishii, Y., Danbara, H. & Okada, N. Genome-wide identification of novel genomic islands that contribute to Salmonella virulence in mouse systemic infection. *FEMS Microbiol. Lett.* **297**, 241–249 (2009).

16. Blanc-Potard, A. B. & Groisman, E. A. The Salmonella selC locus contains a pathogenicity island mediating intramacrophage survival. *EMBO J.* **16**, 5376–5385 (1997).

17. Blanc-Potard, A. B., Solomon, F., Kayser, J. & Groisman, E. A. The SPI-3 pathogenicity island of Salmonella enterica. *J. Bacteriol.* **181**, 998–1004 (1999).

18. Morgan, E., Bowen, A. J., Carnell, S. C., Wallis, T. S. & Stevens, M. P. SiiE is secreted by the Salmonella enterica serovar Typhimurium pathogenicity island 4-encoded secretion system and contributes to intestinal colonization in cattle. *Infect. Immun.* **75**, 1524–1533 (2007).

19. Lorkowski, M., Felipe-López, A., Danzer, C. A., Hansmeier, N. & Hensel, M. Salmonella enterica invasion of polarized epithelial cells is a highly cooperative effort. *Infect. Immun.* **82**, 2657–2667 (2014).

20. Hoiseth, S. K. & Stocker, B. A. Aromatic-dependent Salmonella typhimurium are non-virulent and effective as live vaccines. *Nature* **291**, 238–239 (1981).

21. Hautefort, I., Proença, M. J. & Hinton, J. C. D. Single-Copy Green Fluorescent Protein Gene Fusions Allow Accurate Measurement of Salmonella Gene Expression In Vitro and during Infection of Mammalian Cells. *Appl. Environ. Microbiol.* **69**, 7480–7491 (2003).

22. Datsenko, K. A. & Wanner, B. L. One-step inactivation of chromosomal genes in Escherichia coli K-12 using PCR products. *Proc. Natl. Acad. Sci.* **97**, 6640–6645 (2000).
